# Reconstruction of plant–pollinator networks from observational data

**DOI:** 10.1101/754077

**Authors:** Jean-Gabriel Young, Fernanda S. Valdovinos, M. E. J. Newman

**Affiliations:** Department of Computer Science, University of Vermont, Burlington, Vermont, USA; Vermont Complex Systems Center, University of Vermont, Burlington, Vermont, USA; Center for the Study of Complex Systems, University of Michigan, Ann Arbor, Michigan, USA; Department of Environmental Science and Policy, University of California, Davis, California, USA; Department of Ecology and Evolutionary Biology, University of Michigan, Ann Arbor, Michigan, USA; Department of Physics, University of Michigan, Ann Arbor, Michigan, USA

## Abstract

Empirical measurements of ecological networks such as food webs and mutualistic networks are often rich in structure but also noisy and error-prone, particularly for rare species for which observations are sparse. Focusing on the case of plant-pollinator networks, we here describe a Bayesian statistical technique that allows us to make accurate estimates of network structure and ecological metrics from such noisy observational data. Our method yields not only estimates of these quantities, but also estimates of their statistical errors, paving the way for principled statistical analyses of ecological variables and outcomes. We demonstrate the use of the method with an application to previously published data on plant-pollinator networks in the Seychelles archipelago, calculating estimates of network structure, network nestedness, and other characteristics.

## INTRODUCTION

Network-based methods of analysis have contributed substantially to our understanding of ecological systems by helping us identify structure in the patterns of interaction between species [1–4]. Theoretical studies have shown that such patterns affect the dynamics and stability of ecosystems [5–7]. This is particularly the case for mutualistic networks such as plant-pollinator interactions—our focus in this paper—whose functions are critical to terrestrial biodiversity [8–10] and crop production [10–12].

A central prerequisite for quantitative analysis of network structure and function is accurate network data, and significant effort has been invested in recent years in data gathering for ecological networks of many kinds, including mutualistic networks. There is, however, some debate over whether the observed structure of mutualistic networks represents the true interaction patterns produced by evolutionary and ecological mechanisms, at least to a good approximation [4, 6, 13], or whether, conversely, it is biased by incomplete sampling [14], for instance failing to detect the interactions of rare species [15–18]. In this paper we describe a new technique that aims to give quantitative answers to these questions by applying methods of Bayesian inference to ecological network data. Treating the case of plant-pollinator networks, we show that it is possible to accurately infer interaction network structure from observational data while taking into account confounding variables such as varying species abundances. The output of our calculations is an estimate of the true structure of the network and also a quantification of our uncertainty about this structure. Standard techniques from statistical network science [19, 20] and network ecology [18] can then help us make precise statements about the accuracy of any further conclusions we draw from the network structure. Estimates of interaction certainty can also help us identify interactions that would benefit from greater sampling effort.

The structure of mutualistic networks is typified by several characteristic features [21]: moderate connectance, meaning that a modest fraction of all potential interactions are realized; long-tailed degree distributions, meaning that there are many specialist species with a small number of interactions and a few generalist species with many interactions; and nestedness, meaning that the interactions of the least-connected species are often subsets of the interactions of better-connected species. (These features are not necessarily independent. For instance, it has been suggested that nestedness is itself a consequence of the long-tailed degree-distribution [22].) A significant volume of research has been devoted to explaining these features in terms of ecological and evolutionary mechanisms—see Bascompte and Jordano [6] and Vázquez *et al.* [13] for reviews. Other work, however, has suggested that they can also be generated merely as artifacts of skewed abundance distributions and incomplete sampling, both very common in ecological systems [15, 16]. In particular, Blüthgen *et al.* [15] have shown that nestedness and broad degree distributions can be a result of failure to observe interactions between rare species because of low sampling effort and/or the infrequency of the interactions in question. Findings like this have stimulated further investigations of the effects of sampling bias on network structure [4], both empirically by varying sampling effort in the field [23–27] and theoretically using models of network structure [15, 28–30]. These studies suggest that incomplete sampling strongly underestimates the number of interactions in networks and overestimates the degree of specialization. The approach described in this paper offers one way to address these shortcomings and obtain reliable estimates of the structure of mutualistic networks, free of measurement bias.

The paper is organized as follows. We first outline a first-principles statistical model of plant-pollinator interactions and show how it can be used to estimate network structure from error-prone observational data. Then, we demonstrate these methods with an application to a typical plant-pollinator data set, showing how they give us not only the network structure itself but also statistically principled estimates of quantities such as nestedness. Finally, we give some conclusions and directions for future work.

## RESULTS

### Network reconstruction from observational data

The typical field study of plant-pollinator interactions involves recording instances of potential pollinators (such as insects) visiting plants within a prescribed observation area and over a prescribed period of time. We will refer to these records as *visitation data*. Network ecologists analyze visitation data by constructing networks of plant and pollinator species, where a connection between two species indicates that a plant-pollinator interaction exists between them.

However, the meaning of edges in ecological networks is not always clear [31]. One popular way to transform visitation data into networks is to connect two species when they interact “enough”—say when a pollinator species is seen on the reproductive organ of a plant species a specified number of times—but in this case the precise meaning of an edge will depend on the details of the data collection and the choices made in the analysis. How many visits do we take as evidence of a plant-pollinator interaction? A single visit is probably not enough—it might well be an error or misobservation. Is two enough, or ten, or a hundred? And what about observations that were missed entirely? Other methods of analysis transform the data in different ways, for instance encoding them as weighted networks, possibly with some statistical processing along the way [32]. Even in this case, however, the edges still just count numbers of visits (perhaps transformed in some way), so the resulting networks are effectively histograms in disguise, recording only potential interactions rather than true biological connections.

A more principled approach to network construction begins with a clear definition of what relationship (or relationships) a network’s edges encode [33]. We argue that network ecology often calls for a network of preferred interactions. In the context of plant-pollinator networks the edges of such a network indicate that pollinators preferentially visit certain plant species and they encode a variety of mechanisms that constrain species interactions, such as temporal or spatial uncoupling (i.e., species that do not co-occur in either time or space), constraints due to trait mismatches (e.g., proboscis size very different from corolla size), and physiological-biochemical constraints that prevent the interactions (e.g., chemical barriers). (One can regard preferred interactions as being the opposite of the “forbidden links” described in [34–36]). Preferred interactions are arguably the relevant ones for instance when analyzing the reaction of a network to abrupt changes: when one removes a plant species from a system, for example, the pollinators that prefer it will have to modify their behavior [7, 37, 38]. The interactions we consider are binary—either a species prefers another species or it doesn’t—so the network does not encode varying strengths of interaction.

While the data gathered in a typical field study are certainly reflective of preferred interactions, they are, for many reasons, not perfect measurements of networks of preferred interactions [13, 17]. First, there may be observational errors. While the observers performing the work are usually highly trained individuals, they may nonetheless make mistakes. They may confuse one species for another, which is particularly easy to do for small-bodied insects, or smaller species may be overlooked altogether. Observers may make correct observations but record them wrongly. And there will be statistical fluctuations in the number of visits of an insect species to a plant species over any finite time. For rare interactions there may even be no visits at all if we are unlucky. The insects themselves may also appear to make “mistakes” by visiting plants that they typically do not pollinate. These and other factors mean that the record of observed visits is an inherently untrustworthy guide to the true structure of the network of preferred interactions. Here we develop a statistical method for making estimates of network structure despite these limitations of the data.

#### Model of plant-pollinator data

Consider a typical plant-pollinator study in which some number ***n_p_*** of plant species, labeled by ***i* = 1 *… n_p_***, and some number ***n_a_*** of animal pollinator species, labeled by *j* = 1 *… n_a_*, are under observation for a set amount of time, producing a record of observed visits such that ***M_i j_*** is the number of times plant species ***i*** is visited by pollinator species *j*. Collectively the ***M_i j_*** can be regarded as a data matrix ***M*** with ***n_p_*** rows and ***n_a_*** columns. This is the input to our calculation.

The unknown quantity, the thing we would like to understand, is the network of plant-pollinator interactions. We can think of this network as composed of two sets of nodes, one representing plant species and the other pollinator species, with connections or edges joining each pollinator to the plants it pollinates. In the language of network science this is a bipartite network, meaning that edges run only between nodes of unlike kinds—plants and pollinators—and never between two plants or two pollinators. Such a network can be represented by a second matrix ***B***, called the incidence matrix, with the same size as the data matrix and elements ***B_i j_* = 1** if plant *i* is preferentially visited by pollinator *j* and 0 otherwise.

The question we would like to answer is this: What is the best guess at the structure of the network, represented by ***B***, given the data ***M*?** It is not straightforward to answer this question directly, but it is relatively easy to answer the reverse question. If we imagine that we know ***B***, then we can say what the probability is that we make a specific set of observations ***M***. And if we can do this then the methods of Bayesian inference allow us to invert the calculation and compute ***B*** from a knowledge of ***M*** and hence achieve our goal. The procedure is as follows.

Consider a specific plant-pollinator species pair ***i, j***. How many times do we expect to see ***j*** visit ***i*** if there is, or is not, a preferred interaction between ***i*** and ***j*?** The answer will depend on several factors. First, and most obviously, we expect the number of visits to be higher if ***j*** is in fact a pollinator of *i*. That is, we expect ***M_i j_*** to be larger if ***B_i j_* = 1** than if ***B_i j_* = 0**. Second, we expect there to be more visits if there is greater *sampling effort*—for instance if the period of observation is longer or if the land area over which observations take place is larger [15, 16, 26, 27]. Third, we expect to see more visits for more abundant plant and pollinator species than for less abundant ones, as demonstrated by several studies [28, 30]. And fourth, as discussed above, we expect there to be some random variation in the number of visits, driven by fluctuations in individual behavior and the environment. These are the primary features that we incorporate into our model. It is possible to add others to handle specific situations (see Ref. [39] and the Methods), but we focus on these four here.

We translate these factors into a mathematical model of plant-pollinator interaction as follows. The random variations in the numbers of visits will follow a Poisson distribution for each plant-pollinator pair ***i, j***, parameterized by a single number, the distribution mean ***μ_i j_***, provided only that measurements are made sufficiently far apart to be independent (which under normal conditions they will be). We expect ***μ_i j_*** to depend on the factors discussed above and we introduce additional parameters to represent this dependence. First we introduce a parameter ***r*** to represent the change in the average number of visits when two species are connected (***B_i j_* = 1**), versus when they are not (***B_i j_* = 0**). We write the factor by which the number of visits is increased as **1 + *r*** with ***r* ≥ 0**, so that ***r* = 0** implies no increase and successively larger values of ***r*** give us larger increases. Second, we represent the effect of sampling effort by an overall constant ***C*** that multiplies the mean ***μ_i j_***. The same constant is used for all ***i*** and ***j***, since the same sampling effort is devoted to all plant-pollinator pairs. Third, we assume that the number of visits is proportional to the abundance of the relevant plant and pollinator species: twice as many pollinators of species ***j***, for instance, will mean twice as many visits by that species, and similarly for the abundance of the plant species [13]. Thus the number of visits will be proportional to *σ_i_ τ_j_*, for some parameters ***σ_i_*** and ***τ_j_*** representing the abundances of plant ***i*** and pollinator ***j***, respectively, in suitable units (which we will determine shortly).

Putting everything together, the mean number of observed visits to plant ***i*** by pollinator ***j*** is

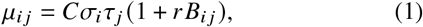

and the probability of observing exactly ***M_i j_*** visits is drawn from a Poisson distribution with this mean:

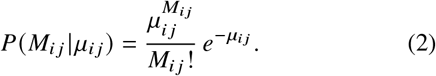

This equation gives us the probability distribution of a single element ***M_i j_*** of the data matrix. Then, combining Eqs. (1) and (2), the data likelihood—the probability of the complete data matrix *M*—is given by the product over all species thus:

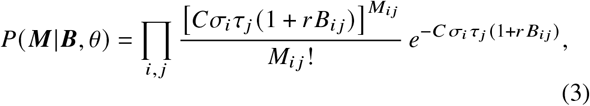

where ***θ*** is a shorthand collectively denoting all the parameters of the model: ***C, r, σ*** and ***τ***. Our model is thus effectively a model of an entire network, rather than single interactions, in contrast with other recent approaches to the modeling of network data reliability [17, 18, 32].

There are two important details to note about this model. First, the definition in **Eq.** (1) does not completely determine ***C*, *σ***, and ***τ*** because we can increase (or decrease) any of these parameters by a constant factor without changing the resulting value of ***μ_i j_*** if we simultaneously decrease (or increase) one or both of the others. In the language of statistics we say that the parameters are not “identifiable.” We can rectify this problem by fixing the normalization of the parameters in any convenient fashion. Here we do this by stipulating that ***σ_i_*** and ***τ_j_*** sum to one, thus:

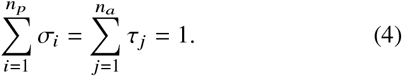

In effect, this makes ***σ_i_*** and ***τ_j_*** measures of relative abundance, quantifying the fraction of individual organisms that belong to each species, rather than the total number. (This definition differs from traditional estimates of pollinator abundance that define the abundance of a pollinator species in terms of its number of observed visits.) Second, there may be other species-level effects on the observed number of visits in addition to abundance, such as the propensity for observers to overlook small-bodied pollinators. There is, at least within the data used in this paper, no way to tell these effects from true variation in abundance—no way to tell for example if there are truly fewer individuals of a species or if they are just hard to see and hence less often observed. As a result, the abundance parameters in our model actually capture a combination of effects on observation frequency. This does not affect the accuracy of the model, which works just as well either way, but it does mean that we have to be cautious about interpreting the values of the parameters in terms of actual abundance. This point is discussed further in the applications below.

#### Bayesian reconstruction

The likelihood of **Eq.** (3) tells us the probability of the data ***M*** given the network ***B*** and parameters ***θ***. What we actually want to know is the probability of the network and parameters given the data, which we can calculate by applying Bayes’ rule in the form

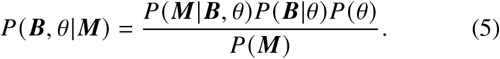

This is the posterior probability that the network has structure ***B*** and parameter values ***θ*** given the observations that were made. There are three important parts to the expression: the likelihood ***P(M|B,θ*)**, the prior probability of the network ***P(B|θ)***, and the prior probability of the parameters ***P(θ)***. The denominator ***P(M)*** we can ignore because it depends on the data alone and will be constant (and hence irrelevant for our calculations) once ***M*** is determined by the observations.

Of the three non-constant parts, the first, the likelihood, we have already discussed—it is given by **Eq.** (3). For the prior on the network ***P*(*B*|*θ***) we make the conservative assumption—in the absence of any knowledge to the contrary—that all edges in the network are a priori equally likely. If we denote the probability of an edge by ***ρ***, then the prior probability on the entire network is

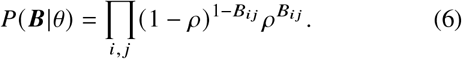

We consider ***ρ*** an additional parameter which is to be inferred from the data and which we will henceforth include, along with our other parameters, in the set ***θ***.

To complete **Eq.** (5), we also need to choose a prior ***P(θ)*** over the parameters. We expect there to be some limit on the value of ***r***, which we impose using a minimally informative prior with finite mean (this distribution turns out to be the exponential distribution). For the remaining parameters we use uniform priors. With these choices, we then have everything we need to compute the posterior probability, **Eq.** (5).

Once we have the posterior probability there are a number of things we can do with it. The simplest is just to maximize it with respect to the unknown quantities ***B*** and ***θ*** to find the most likely structure for the network and the most likely parameter values, given the data. This, however, misses an opportunity for more detailed inference and can moreover give misleading results. In most cases there will be more than one value of ***B*** and ***θ*** with high probability under Eq. (5): there may be a unique maximum of the probability, a most likely value, but there are often many other values that have nearly as high probability and offer plausible network structures competitive with the most likely one. To get the most complete picture of the structure of the network we should consider all these plausible structures.

For example, if all plausible structures are similar to one another in their overall shape then we can be quite confident that this shape is reflective of the true preferred interactions between plant and pollinator species. If plausible structures are widely varying, however, then we have many different candidates for the true structure and our certainty about that structure is correspondingly lower. In other words, by considering the complete set of plausible structures we can not only make an estimate of the network structure but also say how confident we are in that estimate, in effect putting “error bars” on the network.

How do we specify these errors bars in practice? One way is to place posterior probabilities on individual edges in the network. For example, when considering the edge connecting plant ***i*** and pollinator ***j***, we would not ask “**I**s there an edge?” but rather “What is the probability that there is an edge?” Within the formulation outlined above, this probability is given by the average

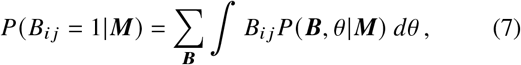

where the sum runs over all possible incidence matrices and the integral over all parameter values. More generally we can compute the average of any function ***f* (*B,θ***) of the matrix ***B*** and/or the parameters ***θ*** thus:

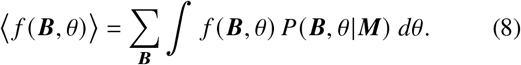

Functions of the matrix and functions of the parameters can both be interesting—the matrix tells us about the structure of the network but the parameters, as we will see, can also reveal important information.

Computing averages of the form (8) is unfortunately not an easy task. A closed-form expression appears out of reach and the brute-force approach of performing the sums and integrals numerically over all possible networks and parameters is computationally intractable in all but the most trivial of cases. The sum over ***B*** alone involves 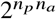 terms, which is normally a very large number.

Instead therefore we use an efficient Monte Carlo sampling technique to approximate the answers. We generate a sample of network/parameter pairs **(*B*_1_*, θ*_1_) *,…, (B_n_, θ_n_*)**, where each pair appears with probability proportional to the posterior distribution of **Eq.** (5). Then we approximate the average of ***f* (*B, θ*)** as

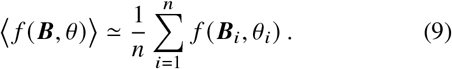

Under very general conditions, this estimate will converge to the true value of the average asymptotically as the number of Monte Carlo samples ***n*** becomes large. Full details of the computations are given in Materials and Methods, and an extensive simulation study of the model is presented in Supplementary Note 1.

#### Checking the model

Inherent in the discussion so far is the assumption that the data can be well represented by our model. In other words, we are assuming there is at least one choice of the network ***B*** and parameters ***θ*** such that the model will generate data similar to what we see in the field. This assumption could be violated if our model is a poor one, but there is nothing in the method described above that would tell us so. To be fully confident in our results we need to be able not only to infer the network structure, but also to check whether that structure is a good match to the data. The Bayesian toolbox comes with a natural procedure for doing this. Given a set of high-probability values of ***B*** and *θ* generated by the method, we can use them in **Eq.** (3) to compute the likelihood ***P(M|B,θ)*** of a data set ***M*** and then sample possible data sets from this probability distribution, in effect recreating data as they would appear if the model were in fact correct. We can then compare these data to the original field data to see if they are similar: if they are then our model has done a good job of capturing the structure in the data.

In the parlance of Bayesian statistics this approach is known as a posterior-predictive assessment [40]. It amounts to calculating the probability

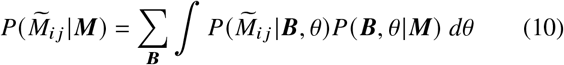

that pollinator species *j* makes 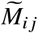 visits to plant species *i* in artificial data sets generated by the model, averaged over many sets of values of ***B*** and ***θ***. We can then use this probability to calculate the average value of 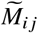 thus:

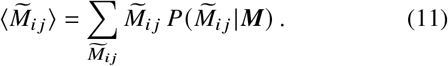

The averages for all plant-pollinator pairs can be thought of as the elements of a matrix 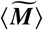, which we can then compare to the actual data matrix ***M***, or alternatively we can calculate a residue 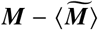. If 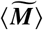 and ***M*** are approximately equal, or equivalently if the residue is small, then we consider the model a good one.

To quantify the level of agreement between the fit and the data we can also compute the discrepancy [40] between the artificial data and ***M*** as

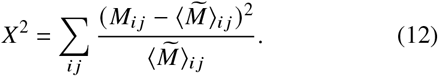

Under the hypothesis that the model is correct, ***X*^2^** follows a chi-squared distribution with ***n_p_ × n_a_*** degrees of freedom [40]. A good fit between model and data is signified by a value of ***X*^2^** that is much smaller than its expectation value of **n_p_ × n_a_**. Note that the calculation of 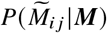 in **Eq.** (10) is of the same form as the one in **Eq.** (8), with 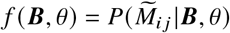, which means we can calculate 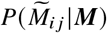 in the same way we calculate other average quantities, using Monte Carlo sampling and **Eq.** (9).

### Application to visitation data sets

#### Well-sampled data

To demonstrate how the method works in practice, we first consider a large data set of plant-pollinator interactions gathered by Kaiser-Bunbury and collaborators [41] at a set of study sites on the island of Mahé in the Seychelles. The data describe the interactions of plant and pollinator species observed over a period of eight months across eight different sites on the island. The data also include measurements of floral abundances for all observation periods and all sites. Our method for inferring network structure does not make use of the abundance measurements, but we discuss them briefly at the end of this section.

The study by Kaiser-Bunbury *et al.* focused particularly on the role of exotic plant species in the ecosystem and on whether restoring a site by removing exotic species would significantly impact the resilience and function of the plant-pollinator network. To help address these questions, half of the sites in the study were restored in this way while the rest were left unrestored as a control group.

As an illustration of our method we apply it to data from one of the restored sites, as observed over the course of a single month in December 2012 (the smallest time interval for which data were available). We pick the site named “Trois-Frères” because it is relatively small but also well sampled. Our calculation then proceeds as shown in Fig. 1. There were 8 plant and 21 pollinator species observed at the site during the month, giving us an 8 21 data matrix ***M*** as shown in Fig. 1a. (Following common convention, the plots of matrices in this paper are drawn with rows and columns ordered by decreasing numbers of observed interactions, so that the largest elements of the data matrix—the darkest squares—are in the top and left of the plot.)

**FIG. 1.**
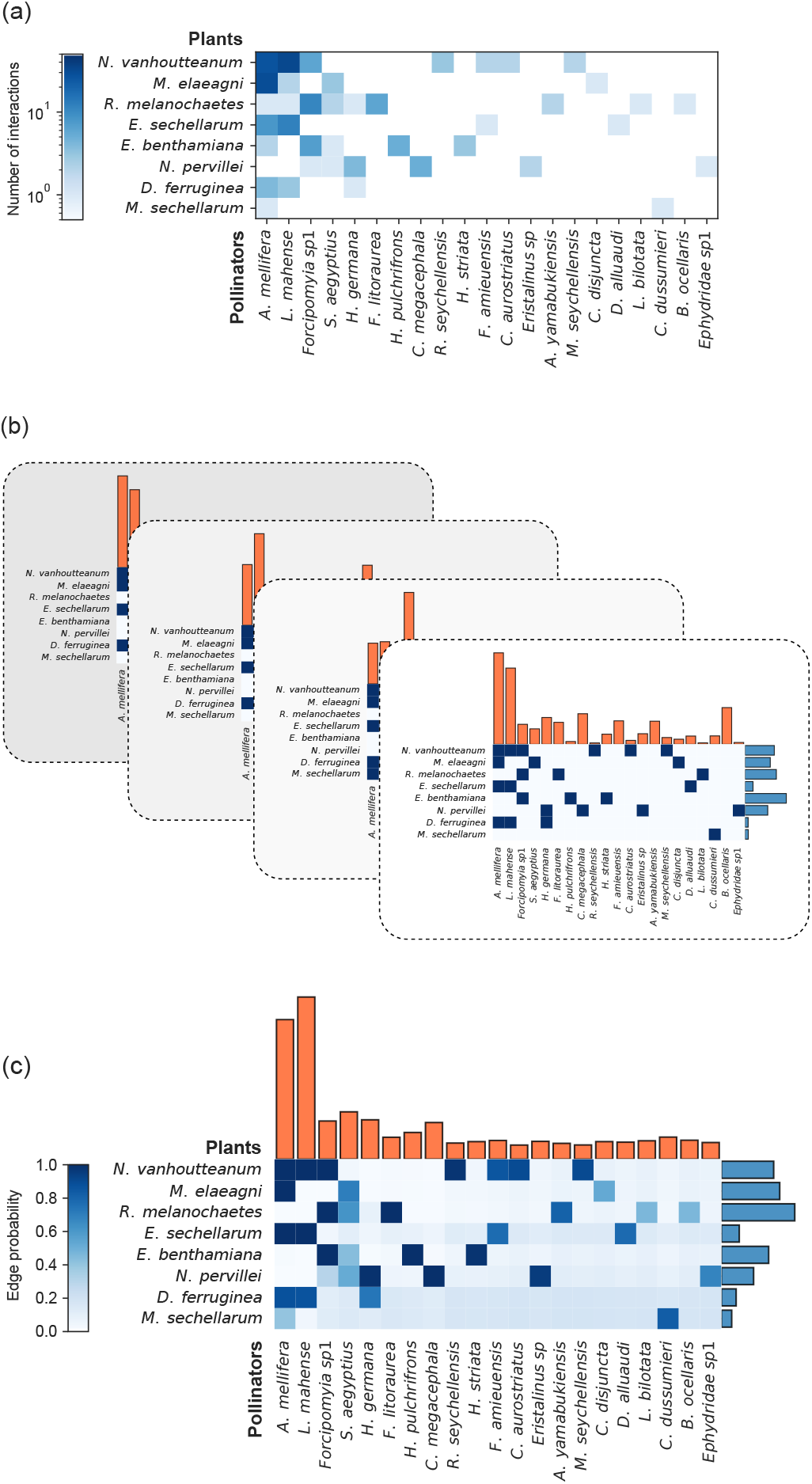
Illustration of the method of this paper applied to data from the study of Kaiser-Bunbury *et al.* [41]. (*a*) We start with a data matrix ***M*** that records the number of interactions between each plant species and pollinator species. Species pairs that are never observed to interact **(*Mi j* = 0)** are shown in white. (*b*) We then draw 2000 samples from the distribution of Eq. (5), four of which are shown in the figure. Each sample consists of a binary incidence matrix *B*, values for the relative abundances ***σ*** and ***τ*** (shown as the orange and blue bar plots, respectively), and values for the parameters *C*, *r*, and *ρ* (not shown). (*c*) We combine the samples using Eqs. (7)-(9) to give an estimate of the probability of each edge in the network and the complete parameter set *θ*. For the data set studied here our estimates of the expected values of the parameters *C*, ***r***, and ***ρ*** are 〈*C*〉 = 20.2, 〈*r*〉 = 45.9, and 〈*ρ*〉 = 0.244.

Now we use our Monte Carlo procedure to draw 2000 sets of incidence matrices ***B*** and parameters *θ* from the posterior distribution of Eq. (5) (Fig. 1b). These samples vary in their structure: some edges, like the one connecting the plant ***N.** vanhoutteanum* and the pollinator ***A.** mellifera*, are present in nearly all samples, while others, like the one between ***M.** sechellarum* and ***A.** mellifera*, appear only a small fraction of the time. Some others never occur at all. Averaging over these sampled networks we can estimate the probability, Eq. (7), that each connection exists in the network of preferred interactions between plant and animal species—see Fig. 1c. Some connections have high probability, close to 1, meaning that we have a high degree of confidence that they exist. Others have probability close to 0, meaning we have a high degree of confidence that they do not exist. And some have intermediate probabilities, meaning we are uncertain about them (such as the ***M.** sechellarum*-***A.** mellifera* connection, which has probability around 0.45). In the latter case the method is telling us that the data are not sufficient to reach a firm conclusion about these connections. Indeed, if we compare with the original data matrix ***M*** in Fig. 1a, we find that most of the uncertain connections are ones for which we have very few observations, relative to the total number of observations for these species—say ***M_i j_* = 1 or 2** for species with dozens of total observations overall.

As we have mentioned, we also need to check whether the model is a good fit to the data by performing a posterior-predictive test. Figure 2 shows the results of this test. The main plot in the figure compares the values of the 40 largest elements of the original data matrix ***M*** with the corresponding elements of the generated matrix 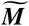. In each case, the original value is well within one standard deviation of the average value generated by the test, confirming the accuracy of the model. The inset of the figure shows the residue matrix 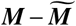, which reveals no systematic bias unaccounted for by the model. The discrepancy ***X*^2^** of Eq. (12) takes the value 26.94 in this case, well below the expected value of *n _p_n_a_* = 168, which indicates that the good fit is not a statistical fluke.

**FIG. 2.**
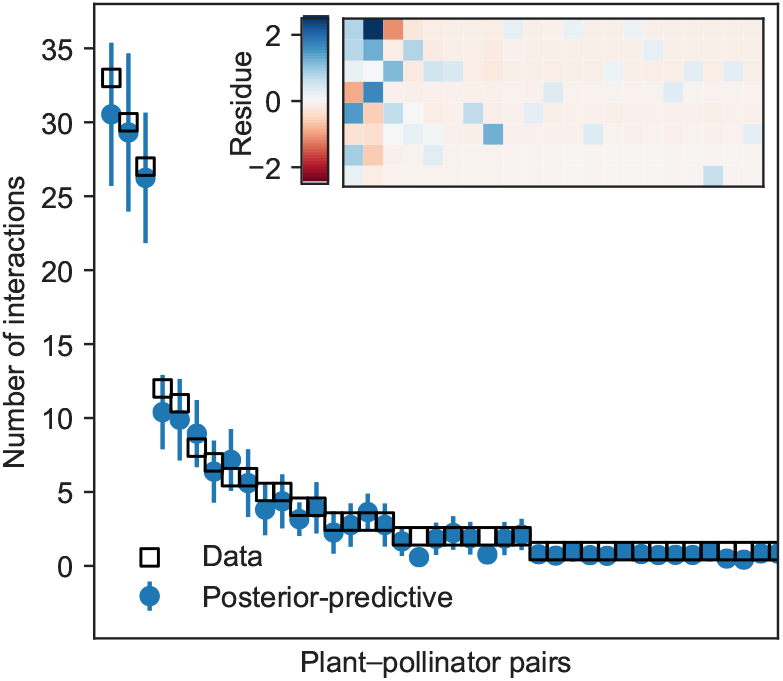
Results of a posterior-predictive test on the data matrix ***M*** for the example data set analyzed in Fig. 1. The main plot shows the error on the 40 largest entries of ***M***, while the inset shows the residue matrix 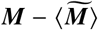. Because the actual data ***M*** are well within one standard deviation of the posterior-predictive mean, the test confirms that the model is a good fit in this case. Error bars correspond to one standard deviation and are computed with *n* = 2000 samples from the posterior distribution.

In addition to inferring the structure of the network itself, our method allows us to estimate many other quantities from the data. There are two primary methods by which we can do this. One is to look at the values of the fitted model parameters, which represent quantities such as the preference ***r*** and species abundances ***σ, τ***. The other is to compute averages of quantities that depend on the network structure or the parameters (or both) from Eq. (9).

As an example of the former approach, consider the parameter *ρ*, which represents the average probability of an edge, also known as the connectance of the network. Figure 3a shows the distribution of values of this quantity over our set of Monte Carlo samples, and neatly summarizes our overall certainty about the presence or absence of edges. If we were certain about all edges in the network, then *ρ* would take only a single value and the distribution would be narrowly peaked. The distribution we observe, however, is somewhat broadened, indicating significant uncertainty. The most likely value of *ρ*, the peak of the distribution, turns out to be quite close to the value one would arrive at if one were simply to assume that every pair of species that interacts even once is connected in the network. This does not mean, however, that one could make this assumption and get good results. As we show below, the network one would derive by doing so would be badly in error in other ways.

**FIG. 3.**
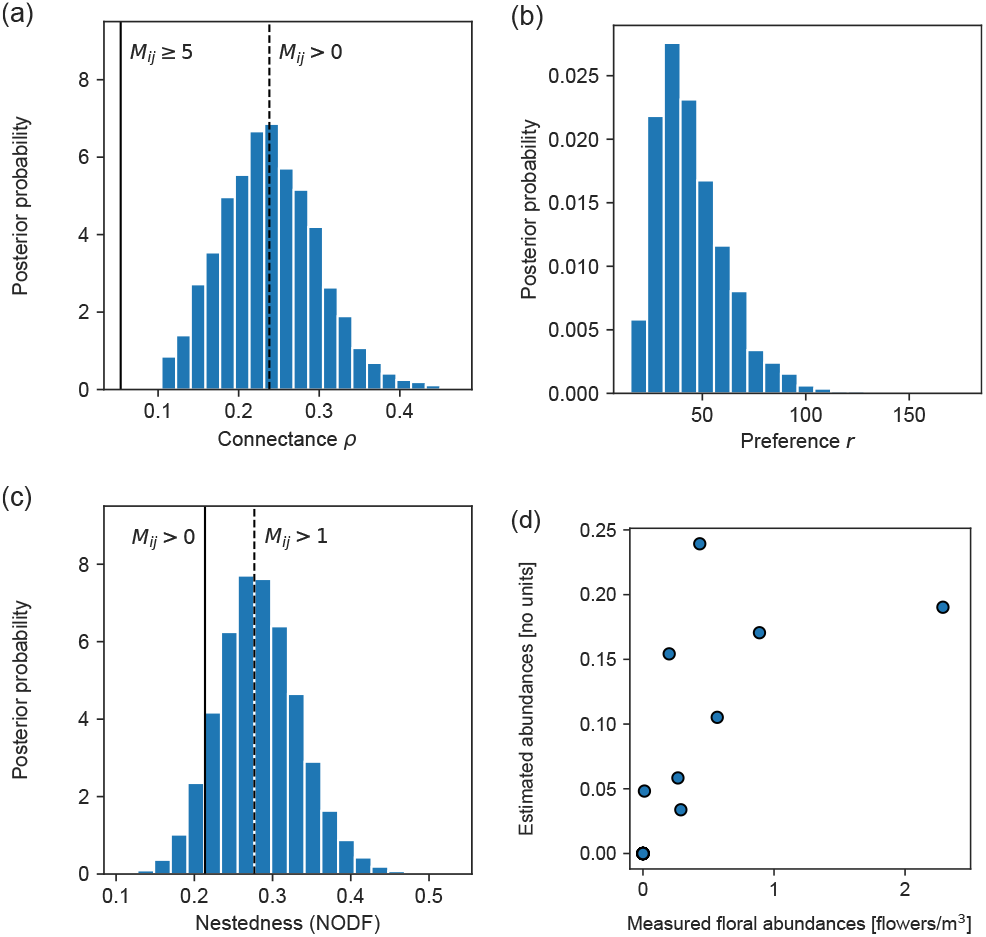
Analyses that can be performed using samples from the posterior distribution of Eq. (5). (*a*) Distribution of the connectance *ρ*. Connectance values for binary networks obtained by thresholding the data matrix at ***M_ij_* > 0** and ***M_ij_* 5** are shown as vertical lines for reference. (*b*) Distribution of the preference parameter *r*. The mean value of *r* is 〈*r*〉 = 45.9 and its mode close to 40, but individual values as high as 100 are possible. (*c*) Distribution of the nestedness measure NODF. Values obtained by thresholding the data matrix at ***M_ij_ >* 0** and ***M_ij_ >* 1** are shown for reference. (*d*) Measured and estimated abundances for each of the plant species **(*R*^2^ = 0.54)**.

Figure 3b shows the distribution of another of the model parameters, the parameter *r*, which measures the extent to which pollinators prefer the plants they normally pollinate over the ones they do not. For this particular data set the most likely value of *r* is around 40, meaning that pollinators visit their preferred plant species about 40 times more often than non-preferred ones, all other things being equal, an impressive level of selectivity on the part of the pollinators.

For the calculation of more complicated network properties we can perform an average over the value of any function ***f(B, θ)***, as long as there is an algorithm to compute it. As an example, Fig. 3c shows a calculation of the quantity known as “Nestedness based on Overlap and Decreasing Fill” **(NODF)**, a measure of the nestedness property discussed in the introduction. This quantity measures the extent to which specialist species—those with relatively few interactions—tend to interact with a subset of the partners of generalist species [42]. While it is complicated to compute NODF analytically, due to the fact that one must order the species by degrees [22], it is straightforward to calculate it within our framework: we simply calculate the value for each sampled network ***B*** and plot the resulting distribution. Interestingly, the most likely value of **NODF** is significantly different from the one we would calculate had we assumed, as discussed above, that a single interaction is sufficient to consider two species connected. On the contrary, we find that the system is almost certainly more nested than this simple analysis would conclude.

In Fig. 3d, we compare the values of our estimated floral abundance parameters *σ* to the measured abundances reported by Kaiser-Bunbury *et al.* [41]. These parameters are not measures of abundance in the usual sense, because they combine actual abundance (quantity or density) with other characteristics such as ease of observation. We do find a correlation between the estimated and observed abundances, but it is relatively weak **(*R*^2^ = 0.54)**, signaling significant disagreement, on which we elaborate in the discussion section.

#### Undersampled data

As we have pointed out, the connections in the network about which we are most uncertain tend to be ones that are *undersampled*, i.e., those for which we have only a small amount of data. In an ideal world we could address this problem by taking more data, but it is rare that we have the opportunity to do this. More commonly the data have already been gathered and our task is to produce the best results we can with those data. There are nonetheless some remedies open to us, such as aggregating data over different geographical areas or time windows. In Fig. 4 we compare the edge probabilities estimated from data recorded individually at the four “restored” sites in the Mahé study during October 2012 to the edge probabilities we obtain when we aggregate these observations into a single data matrix and only then estimate the network. (We use restored sites observed during the same month because they are likely to be ecologically similar, meaning the data are measuring approximately the same system.) Comparison of the two distributions shows—as we would hope—that there are fewer uncertain edges in the aggregated network than in its disaggregated parts, i.e., there are fewer edges with probabilities in the middle of the distribution and more with probabilities close to zero or one.

**FIG. 4.**
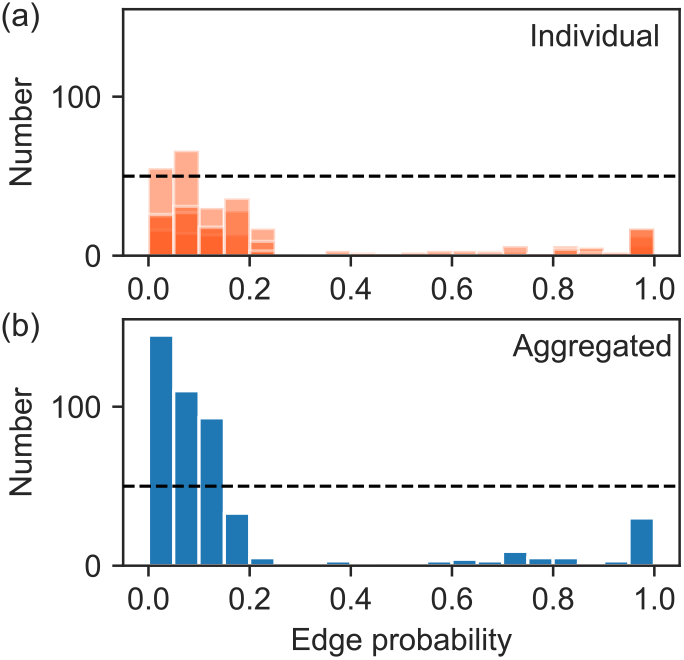
Illustration of the effect of data aggregation on edge uncertainty. (*a*) Histogram of the edge probabilities ***P(Bi j = 1|M)*** for the four restored sites in the Mahé study as observed in October 2012 and analyzed individually. (*b*) Equivalent histogram after aggregating the data over the sites and then estimating a single network from the resulting data matrix. The horizontal lines, both drawn at fifty observations—are added merely as a guide to the eye. Note how the upper histogram has more mass near the middle of the plot, while the lower one has most of its mass close to probability zero or one, indicating greater certainty in the positions of the edges in the aggregated data.

In other cases neither aggregation nor gathering more data is possible, for instance when reanalyzing a data set already collected by others or already maximally aggregated. Such data sets record the results of observational studies that are already over, and may contain too few observations, but our approach still allows us to perform rigorous inference in these circumstances.

For instance, Jordano *et al.* [44] used dozens of existing plant-pollinator and plant-frugivore data sets to argue that the degree distributions of mutualistic networks have a long tail, but this conclusion is undermined by issues with undersampling. As an example, one of the data sets they studied, originally gathered by Inouye and Pyke [43], records 1314 individual interactions over a period of 3 months in Kosciusko National Park, Australia, between 40 plants and 85 pollinator species, which works out to an average of 0.386 unique observations per species pair. Is this sampling effort sufficient to establish edges with certainty? As a point of reference, the data analyzed in Fig. 1 comprises 201 observations between 8 plants and 21 pollinators species for an average of 1.196 observations per pair of species, and the aggregated data of Fig. 4 contain 1.420 observations for every pair. Nonetheless, there is uncertainty about some of the connections in these reconstructed networks; this suggests that the network of Inouye and Pyke, with less than a third as much data per species pair, will contain significant uncertainty.

Even so, our method allows us to make inferences about this network. In Fig. 5, we show estimates of the degree distributions of both plant and pollinator nodes in the network obtained from the posterior distribution ***P(B|M)***, along with naive estimates calculated by thresholding the (undersampled) data as in the study by Jordano *et al.* [44]. As the figure shows, the results derived from the two approaches are very different. The thresholded degree distributions were classified as scale-free by Jordano *et al.*, but this classification no longer holds once we account for the issues with the data; the inferred degree distributions are in this case well-modeled as Poisson distributions of means 5.53 and 2.60 for plants and pollinators respectively and the power-law form is a poor fit. On the other hand, the abundance parameters of the model, shown in Fig. 5, do appear to have a broad distribution, an interesting finding that calls for a rethinking of the relationship between abundances and degree distributions. It is generally thought that interactions will tend to be evenly distributed under an even distribution of abundance [13] but here the opposite seems to be true.

**FIG. 5.**
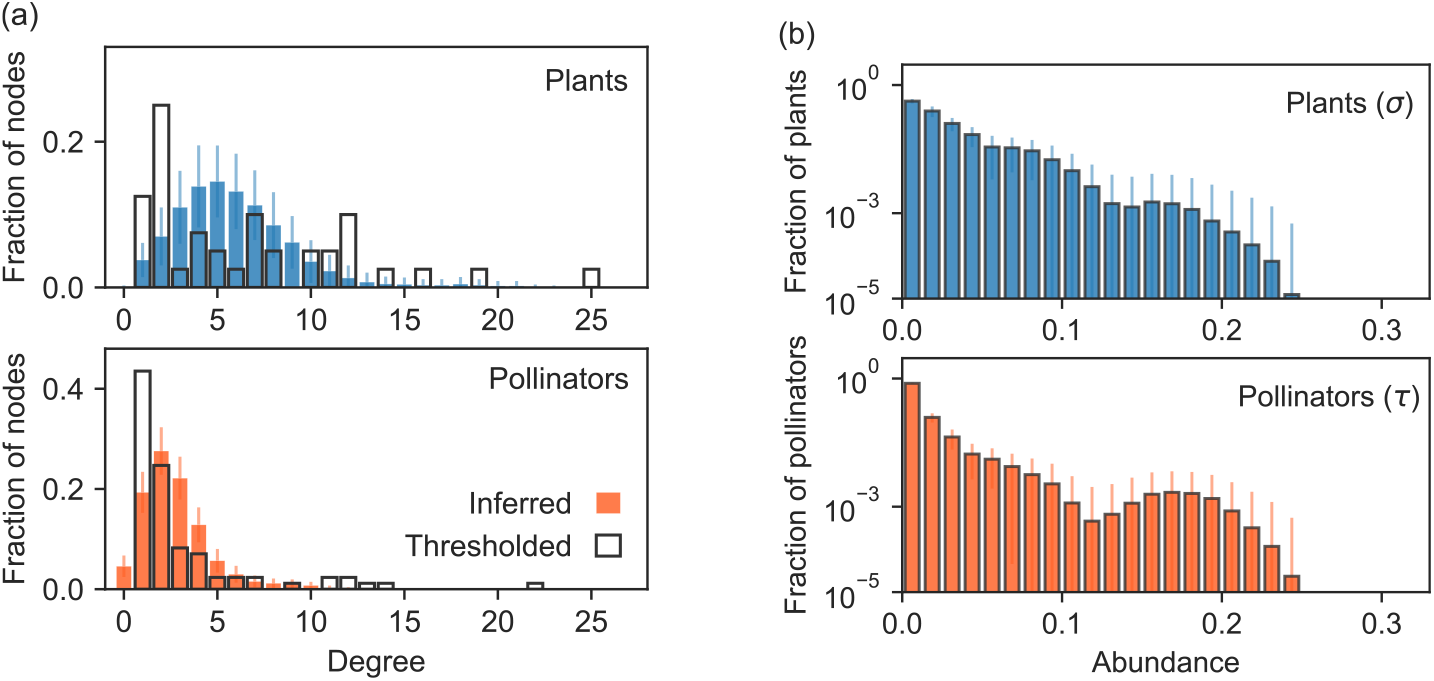
Distributions of species-level parameters for a network of plants and pollinators in Kosciusko National Park, Australia, from the study by Inouye and Pyke [43]. (*a*) Thresholded degree distributions calculated by connecting species *i* and *j* with an edge if ***Mij > 0***. Inferred degree distributions are calculated using the method of this paper, averaging the fraction *pk* of nodes with a given degree *k* over *n* = 2000 Monte Carlo samples. (*b*) Inferred distributions of abundances ***σ*** and ***τ***, calculated as a histogram over ***n* = 2000** Monte Carlo samples of the abundance parameters of the fitted model. Error bars correspond to one standard deviation in all cases.

## DISCUSSION

In this paper, we have proposed a statistical model of plant-pollinator interactions and shown how it can be used to infer the structure and properties of empirical plant-pollinator networks from noisy, error-prone measurements. The model employs elementary ecological insights to create an expressive and versatile structure that can capture the pattern of interactions in a wide range of ecosystems. We use the toolbox of Bayesian statistics to develop both an inference algorithm and a model checking procedure for the model. Our methods explicitly allow for the possibility that there are multiple plausible networks that could fit a given set of observations, a hallmark of Bayesian analysis. Doing this allows us to make accurate deductions even in cases where data sets are small and the number of model parameters is large.

Applying our method to previously published plant-pollinator visitation data, we arrive at a number of conclusions. First, our analysis confirms previous findings that there is some uncertainty in measured values of the connectance [23–27] and that moderate connectance [6] seems to hold in plant-pollinator networks even once we account for uncertainty (Fig. 3a). Second, we have found that pollinators strongly prefer the plants they normally visit over ones they do not, with pollinators visiting their preferred plant species about 40 times more often than non-preferred ones in our results (Fig. 3b). This highlights the strong selectivity of pollinators for the plant species they usually visit. Third, we have found that networks reconstructed using our method are more nested than networks built using thresholds of one or a few visits to determine plant-pollinator interactions, which supports the longstanding claim that plant-pollinator networks are nested [6]. Finally, our analysis suggests that the distribution of number of interactions of a species (the degree distribution) is less skewed than previously thought [44]. This result supports recent findings showing that incomplete sampling strongly underestimates the number of interactions and overestimates the degree of specialization.

Our model and inference algorithm also give an estimate of species abundances. As we have argued, these estimates actually capture a combination of effects on observation frequency beyond just plain abundance, which helps to explain why, as we have seen, measured and estimated floral abundances are correlated but not strongly so. Disagreements between measured and estimated abundances were observed previously by Vazquez *et al.* [28], who used null models to show that measured abundances cannot in general explain the form of visitation matrices. Taken together, these results indicate that the frequency of observed interaction between plants and pollinators is not in fact proportional to their plain abundances (defined as quantity or density of individuals), but instead incorporates a range of factors potentially including abundance, ease of observation, network effects, and others [45]. One candidate for a possible additional factor that could play a role is adaptive foraging by pollinators, which has been shown to influence the structure of ecological networks [4, 46]. Adaptive foraging occurs, for example, when pollinators deliberately visit less abundant plants more often if those plants contain more food (such as nectar or pollen) relative to more abundant plants with less food [7]. Our estimated abundance parameters automatically include such factors where traditional field estimates of pollinator abundance—such as the number of visits of a pollinator species—do not. Analyses that use traditional estimates of abundance, as in Refs. [15, 16], may as a result fail to control for significant species-level effects on observed visitation rates [13]. We would therefore argue that best practice calls for the use of estimated abundances like those proposed here rather than traditional ones when estimating networks of preferred interactions.

There are a number of ways in which the approach presented here could be extended. The method as described assumes an ecosystem that is more or less static, but ecosystems can change rapidly with the seasons. One could imagine a dynamic variant of the model that allows parameters to evolve over time, or networks with several levels of preference, allowing for more nuanced description of plant-pollinator systems. On the applications side, we have limited our analysis to the important case of plant-pollinator networks, but similar methods could be applied to other types of ecological networks, allowing us to better separate signal from noise in those domains too.

## METHODS

As outlined in the main text, our method relies on a generative network model in which observed visits to plants by pollinators are considered noisy measurements of an unobserved underlying plant-pollinator network. This formulation allows us to frame the task of determining the network structure as a Bayesian inference problem [31, 47–49] in which the probability of the network having incidence matrix ***B*** given a data matrix ***M*** is

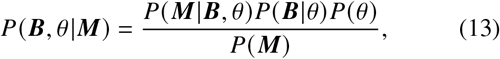

where *θ* are model parameters and ***P(M)*** is an unimportant normalizing constant. The element ***M_i__j_*** of matrix ***M*** records the number of times insects of species ***j*** are seen to pollinate plant species ***i***, while ***B_i j_* = 0, 1** encodes the presence or absence of an edge between the two species in the plant-pollinator network. Both matrices are of dimension *n _p_ n_a_* where ***n_p_*** is the number of plants and ***n_a_*** is the number of pollinators.

We model the number of visits ***M_i j_*** as a Poisson random variable with mean

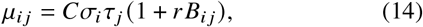

and use independent priors on all parameters with

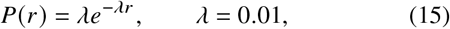

and uniform priors on ***C, σ***, and ***τ***. We further assume that edges are *a priori* equally likely with probability *ρ* and use a uniform prior distribution on *ρ* itself. This leads to

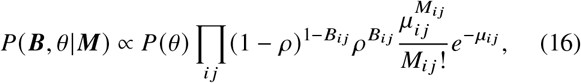

with ***P(θ) ∝ P(r)***. We note that in this Bayesian formulation, one can easily model interaction specific traits [39] or account for known biology like trait-matching [13, 50] by altering the priors on ***B_ij_*** for a particular pair of species ***i, j*** [48].

### Bayesian reconstruction of networks

Given the probability distribution in Eq. (16) there are a number of approaches we could take. Following [47, 48] we could employ an expectation-maximization (EM) algorithm to calculate the distribution over potential network structures and a point estimate of *θ* or, following [49], we could integrate out the parameters *θ* and then sample from the resulting marginal distribution on ***B***. Neither of these approaches is completely satisfactory here however, the first because point estimates of the parameters can be unreliable for large models such as ours, and the second because the values of the model parameters are actually of interest to us, so we would prefer not to eliminate them.

Instead therefore we make use of a technique from the literature on finite mixture models [51] to sample efficiently from the joint distribution of both ***B*** and *θ* and hence estimate both. First, we sample values of the parameters *θ* from their marginal distribution

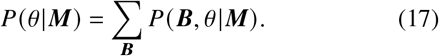

The sum over ***B*** can be carried out analytically because the particular ***(P B,θ|M)*** defined in Eq. (17) can be written in the form

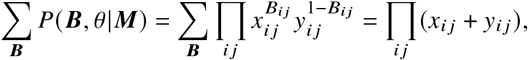

where *x_i j_* and *y_i j_* combine all the terms associated with the situation where there is/is not an edge. We then find that

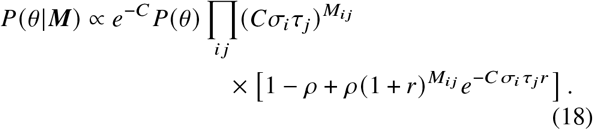

We can now sample from this distribution using standard methods such as Hamiltonian Monte Carlo—see below. This gives us our estimates of the parameter values.

For given values of the parameters we then estimate the network ***B*** itself by sampling from the distribution

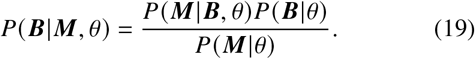

Using the previous expressions for the likelihood ***P*(*M*|*B,θ*)** and ***P*(*B*|*θ*)**—Eqs. (3) and (6) of the Results—and noting that the denominator *P*(*M* |*θ*) is proportional to Eq. (18), we find

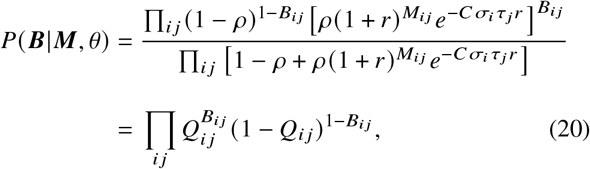

where

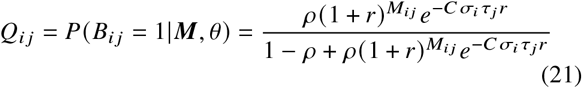

is the posterior probability of an edge between species ***i*** and ***j***, given the parameters ***θ***.

We now simply average ***Q_ij_*** over our sampled values of the parameters ***θ*** to get the expected probability of an edge between any pair of nodes. More generally, we can calculate an estimate of any function ***f(B,θ)*** by drawing *m* samples ***θ_K_*** of the parameter set and *n* random incidence matrices ***B_l_(θ_K_)*** for each set, with edges appearing independently with probabilities ***Q_i j_*** given by (21), then averaging:

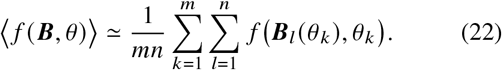

### Implementation

In our implementation of this approach we sample parameters ***θ*** from the distribution of Eq. (18) using the technique known as Hamiltonian Monte Carlo (HMC). In HMC one defines an inertial mechanics in a position space equivalent to the space of the parameters, with auxiliary momenta chosen so that the dynamics under the corresponding Hamilton’s equations samples from the desired distribution [52]. We implement the calculation in Stan, a probabilistic programming language that automatically performs HMC sampling for arbitrary target distributions [53]. In practice, the program operates on the log of the posterior probability, which for our distribution (18) has the form log ***P*(*θ* | *M*) = −*C* + *_i j_* (*X_i j_* + *Y_i j_*)** where

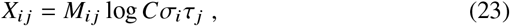

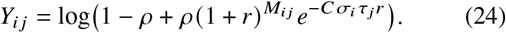

To avoid potential over- or underflow and ensure numerical stability we rewrite the latter expression slightly by defining

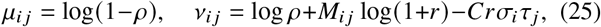

and then writing

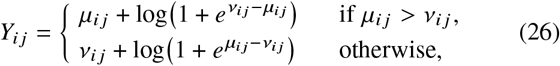

which ensures that *Y_i j_* is always a manageable number.

An important practical consideration is verifying the convergence of the Monte Carlo algorithm. **HMC** mixes rapidly, but, like all Monte Carlo methods, it can sometimes become trapped at local optima. To ensure representative sampling of the posterior distribution, we therefore perform multiple Monte Carlo runs from random initial states and if any of the runs converges to a region of significantly smaller probability than the others then we repeat the entire calculation. In the example calculations given in the paper we perform four runs, with an equilibration period of 5000 Monte Carlo steps each, followed by taking 500 samples.

### Quantifying error using posterior predictive assessment

A crucial part of the model fitting process is assessing whether the model is a good fit to the data. In the main text we argue that a so-called posterior predictive test is a good way of making this assessment. The idea is to generate a new artificial data set 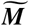 from the model using the values of the model parameters derived from the fit to the input data ***M***. If we find that 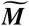 looks similar to the input data then our model has done a good job of capturing the structure of the data.

To carry out this procedure we need to calculate the posterior predictive distribution for species pair ***i, j*** given by

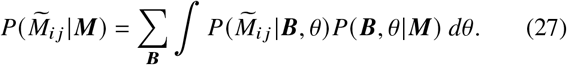

Since the likelihood ***P(M|B, θ)***, Eq. (3), factors into separate terms for each plant-pollinator pair *i, j*, this expression can with a little work be simplified to

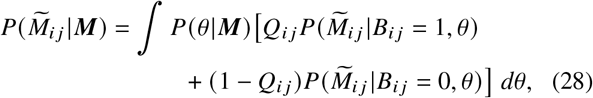

and the integral can then be approximated by simply averaging over the set of sampled values of *θ*.

Two particularly useful statistics for the posterior predictive test are the mean and the variance of 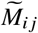, which in this case are equal since 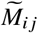 by definition has a Poisson distribution for given ***B*** and *θ*. Both are to a good approximation given by

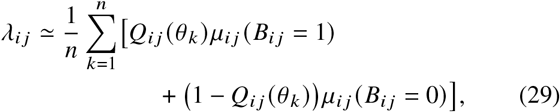

where ***μ_i j_*** is the mean defined in Eq. (14).

### Description of the data sets

The data analyzed in Figs. 1–4 were gathered by Kaiser-Bunbury *et al.* [41] on inselbergs (steep-sided monolithic rocky outcroppings) on the tropical granitic island of Mahé, located in the Indian Ocean. The vegetation on the inselbergs is characterized by short trees, shrubs, and an absence of flowering herbs. The data we analyze includes records of the visits of pollinator species to all plant species found in each of the eight inselbergs, observed between September 2012 and April 2013 during the island’s eight-month-long tropical flowering season. Species visiting flowers were recorded as pollinators if they touched the sexual parts of the flowers within a standard observation window of 30 minutes [54]. Floral abundances were obtained by counting flowers in 1-meter cubes randomly located along transects spanning the inselbergs. The visit data were used to generate 64 data matrices of plant-pollinator interactions, one for each period and location. Our primary analysis focuses on the matrix for the site known as Trois-Frères as observed during the month of December 2012. We chose this data set primarily because it is relatively small and hence easy to visualize.

The data analyzed in Fig. 5 were gathered by Inouye and Pyke [43] in the Kosciusko National Park, Australia, between December 21, 1983, and March 30, 1984. The observations were made in 26 plots of 2m × 2m, chosen before the flowering season and encompassing an alpine zone at elevations ranging from 1940 to 2040 meters and a montane habitat at elevations of 1860 to 1920 meters. Flowers were counted roughly every second day. Insect visitation data were collected through incidental observations made during the phenological censuses of the flowers as well as dedicated observation periods of 10 minutes length, spread throughout the study duration. The data set of Inouye and Pyke [43] is only one of several data sets re-analyzed by Jordano *et al.* [44]. We chose this data set because it is somewhat undersampled, making it a good example of a situation where our method can improve network estimates.

## CODE AVAILABILITY

Reference implementations in `stan` and `python` of the methods described in this study are freely available online [56].

## ACKNOWLEDGMENTS

We thank Alec Kirkley, George Cantwell, and Maria Riolo for helpful discussions. This work was funded in part by the James S. McDonnell Foundation (JGY) and the US National Science Foundation under grants DEB-1834497 (FSV) and DMS-1710848 and DMS-2005899 (MEJN), as well as University of Michigan MICDE grant U061182 (FSV).

## DATA AVAILABILITY

The Mahé visitation data used in this study are available as supplementary material to Kaiser-Bunbury *et al.* [41]. The data of Inouye and Pyke [43] analyzed in Fig. 5 can be downloaded from the Web of Life data base [55], available at http://www.web-of-life.es, under the network identifier M_PL_019.

## Notes

### Competing Interest Statement

The authors have declared no competing interest.

## REFERENCES

1 N. D. Martinez, Artifacts or attributes? Effects of resolution on the Little Rock Lake food web. Ecol. Monogr. 61, 367–392 (1991).

2 J. Bascompte, P. Jordano, C. J. Melian, and J. M. Olesen, The nested assembly of plant-animal mutualistic networks. Proc. Natl. Acad. Sci. U.S.A. 100, 9383–9387 (2003).

3 E. Thébault and C. Fontaine, Stability of ecological communities and the architecture of mutualistic and trophic networks. Science 329, 853–856 (2010).

4 F. S. Valdovinos, Mutualistic networks: Moving closer to a predictive theory. Ecol. Lett. 22, 1517–1534 (2019).

5 U. Brose, R. J. Williams, and N. D. Martinez, Allometric scaling enhances stability in complex food webs. Ecol. Lett. 9, 1228– 1236 (2006).

6 J. Bascompte and P. Jordano, Mutualistic Networks. Princeton University Press, Princeton, NJ (2014).

7 F. S. Valdovinos, B. J. Brosi, H. M. Briggs, P. Moisset de Espanés, R. Ramos-Jiliberto, and N. D. Martinez, Niche partitioning due to adaptive foraging reverses effects of nestedness and connectance on pollination network stability. Ecol. Lett. 19, 1277–1286 (2016).

8 J. N. Thompson, The Coevolutionary Process. University of Chicago Press, Chicago, IL (1994).

9 J. Ollerton, R. Winfree, and S. Tarrant, How many flowering plants are pollinated by animals? Oikos 120, 321–326 (2011).

10 J. Ollerton, Pollinator diversity: Distribution, ecological function, and conservation. Annu. Rev. Ecol. Evol. Syst. 48, 353–376 (2017).

11 S. G. Potts, V. Imperatriz-Fonseca, H. T. Ngo, M. A. Aizen, J. C. Biesmeijer, T. D. Breeze, L. V. Dicks, L. A. Garibaldi, R. Hill, J. Settele, and A. J. Vanbergen, Safeguarding pollinators and their values to human well-being. Nature 540, 220–229 (2016).

12 L. A. Garibaldi, I. Steffan-Dewenter, R. Winfree, M. A. Aizen, R. Bommarco, S. A. Cunningham, C. Kremen, L. G. Carvalheiro, L. D. Harder, O. Afik, et al., Wild pollinators enhance fruit set of crops regardless of honey bee abundance. Science 339, 1608–1611 (2013).

13 D. P. Vázquez, N. Blüthgen, L. Cagnolo, and N. P. Chacoff, Uniting pattern and process in plant-animal mutualistic networks: A review. Ann. Bot. 103, 1445–1457 (2009).

14 R. H. Gibson, B. Knott, T. Eberlein, and J. Memmott, Sampling method influences the structure of plant–pollinator networks. Oikos 120, 822–831 (2011).

15 N. Blüthgen, J. Fründ, D. P. Vázquez, and F. Menzel, What do interaction network metrics tell us about specialization and biological traits? Ecology 89, 3387–3399 (2008).

16 N. Blüthgen, Why network analysis is often disconnected from community ecology: A critique and an ecologist’s guide. Basic Appl. Ecol. 11, 185–195 (2010).

17 A. R. Cirtwill, A. Eklöf, T. Roslin, K. Wootton, and D. Gravel, A quantitative framework for investigating the reliability of empirical network construction. Methods Ecol. Evol. 10(6), 902–911 (2019).

18 T. Poisot, A. R. Cirtwill, K. Cazelles, D. Gravel, M.-J. Fortin, and D. B. Stouffer, The structure of probabilistic networks. Methods Ecol. Evol. 7, 303–312 (2016).

19 P. Parchas, F. Gullo, D. Papadias, and F. Bonchi, Uncertain graph processing through representative instances. ACM Trans. Database Syst. 40, 1–39 (2015).

20 A. Khan, Y. Ye, and L. Chen, On Uncertain Graphs. Morgan & Claypool Publishers, San Rafael, CA (2018).

21 J. Bascompte and P. Jordano, Plant-animal mutualistic networks: The architecture of biodiversity. Annu. Rev. Ecol. Evol. Syst. 38, 567–593 (2007).

22 C. Payrató-Borràs, L. Hernández, and Y. Moreno, Breaking the spell of nestedness: The entropic origin of nestedness in mutualistic systems. Phys. Rev. X 9, 031024 (2019).

23 A. Nielsen and J. Bascompte, Ecological networks, nestedness and sampling effort. J. Ecol. 95, 1134–1141 (2007).

24 T. Petanidou, A. S. Kallimanis, J. Tzanopoulos, S. P. Sgardelis, and J. D. Pantis, Long-term observation of a pollination network: Fluctuation in species and interactions, relative invariance of network structure and implications for estimates of specialization. Ecol. Lett. 11, 564–575 (2008).

25 S. J. Hegland, J. Dunne, A. Nielsen, and J. Memmott, How to monitor ecological communities cost-efficiently: The example of plant-pollinator networks. Biol. Cons. 143, 2092–2101 (2010).

26 N. P. Chacoff, D. P. Vázquez, S. B. Lomáscolo, E. L. Stevani, J. Dorado, and B. Padrón, Evaluating sampling completeness in a desert plant-pollinator network. J. Anim. Ecol. 81, 190–200 (2012).

27 A. Rivera-Hutinel, R. O. Bustamante, V. H. Marín, and R. Medel, Effects of sampling completeness on the structure of plant-pollinator networks. Ecology 93, 1593–1603 (2012).

28 D. P. Vázquez, C. J. Melián, N. M. Williams, N. Blüthgen, B. R. Krasnov, and R. Poulin, Species abundance and asymmetric interaction strength in ecological networks. Oikos 116, 1120– 1127 (2007).

29 I. Bartomeus, Understanding linkage rules in plant-pollinator networks by using hierarchical models that incorporate pollinator detectability and plant traits. PLOS ONE 8, e69200 (2013).

30 J. Fründ, K. S. McCann, and N. M. Williams, Sampling bias is a challenge for quantifying specialization and network structure: Lessons from a quantitative niche model. Oikos 125, 502–513 (2016).

31 I. Brugere, B. Gallagher, and T. Y. Berger-Wolf, Network structure inference, a survey: Motivations, methods, and applications. ACM Comput. Surv. 51, 24:1–24:39 (2018).

32 D. R. Farine and A. Strandburg-Peshkin, Estimating uncertainty and reliability of social network data using bayesian inference. R. Soc. Open Sci. 2, 150367 (2015).

33 C. T. Butts, Revisiting the foundations of network analysis. Science 325, 414–416 (2009).

34 D. P. Vázquez, N. Chacoff, and L. Cagnolo, Evaluating multiple determinants of the structure of plant-animal mutualistic networks. Ecology 90, 2039–2046 (2009).

35 P. Jordano, Patterns of mutualistic interactions in pollination and seed dispersal: connectance, dependence asymmetries, and coevolution. Am. Nat. 129, 657–677 (1987).

36 P. Jordano, Sampling networks of ecological interactions. Funct. Ecol. 30, 1883–1893 (2016).

37 C. N. Kaiser-Bunbury, S. Muff, J. Memmott, C. B. Müller, and A. Caflisch, The robustness of pollination networks to the loss of species and interactions: a quantitative approach incorporating pollinator behaviour. Ecol. Lett. 13, 442–452 (2010).

38 R. Ramos-Jiliberto, F. S. Valdovinos, P. M. de Espanés, and J. D. Flores, Topological plasticity increases robustness of mutualistic networks. J. Anim. Ecol. 81, 896–904 (2012).

39 C. N. Kaiser-Bunbury, D. P. Vázquez, M. Stang, and J. Ghazoul, Determinants of the microstructure of plant–pollinator networks. Ecology 95(12), 3314–3324 (2014).

40 A. Gelman, X.-L. Meng, and H. Stern, Posterior predictive assessment of model fitness via realized discrepancies. Stat. Sin. 6, 733–760 (1996).

41 C. N. Kaiser-Bunbury, J. Mougal, A. E. Whittington, T. Valentin, R. Gabriel, J. M. Olesen, and N. Blüthgen, Ecosystem restoration strengthens pollination network resilience and function. Nature 542, 223–227 (2017).

42 M. Almeida-Neto, P. Guimaraes, P. R. Guimaraes Jr, R. D. Loyola, and W. Ulrich, A consistent metric for nestedness analysis in ecological systems: Reconciling concept and measurement. Oikos 117, 1227–1239 (2008).

43 D. W. Inouye and G. H. Pyke, Pollination biology in the snowy mountains of Australia: comparisons with montane Colorado, USA. Aust. J. Ecol. 13, 191–205 (1988).

44 P. Jordano, J. Bascompte, and J. M. Olesen, Invariant properties in coevolutionary networks of plant-animal interactions. Ecol. Lett. 6, 69–81 (2003).

45 O. Ovaskainen, N. Abrego, P. Halme, and D. Dunson, Using latent variable models to identify large networks of species-to-species associations at different spatial scales. Methods Ecol. Evol. 7, 549–555 (2016).

46 F. S. Valdovinos, R. Ramos-Jiliberto, L. Garay-Narváez, P. Urbani, and J. A. Dunne, Consequences of adaptive behaviour for the structure and dynamics of food webs. Ecol. Lett. 13, 1546– 1559 (2010).

47 M. E. J. Newman, Network structure from rich but noisy data. Nat. Phys. 14, 542–545 (2018).

48 M. E. J. Newman, Estimating network structure from unreliable measurements. Phys. Rev. E 98, 062321 (2018).

49 T. P. Peixoto, Reconstructing networks with unknown and heterogeneous errors. Phys. Rev. X 8, 041011 (2018).

50 M. Pichler, V. Boreux, A. Klein, M. Schleuning, and F. Hartig, Machine learning algorithms to infer trait-matching and predict species interactions in ecological networks. Methods Ecol. Evol. 11, 281–293 (2019).

51 A. Gelman, H. S. Stern, J. B. Carlin, D. B. Dunson, A. Vehtari, and D. B. Rubin, Bayesian Data Analysis. Chapman and Hall/CRC, New York, NY, 3rd edition (2013).

52 M. Betancourt, A conceptual introduction to Hamiltonian Monte Carlo. Preprint arxiv:1701.02434 (2017).

53 B. Carpenter, A. Gelman, M. D. Hoffman, D. Lee, B. Goodrich, M. Betancourt, M. Brubaker, J. Guo, P. Li, and A. Riddell, Stan: A probabilistic programming language. J. Stat. Softw. 76, 1–32 (2017).

54 C. N. Kaiser-Bunbury, T. Valentin, J. Mougal, D. Matatiken, and J. Ghazoul, The tolerance of island plant–pollinator networks to alien plants. J. Ecol. 99, 202–213 (2011).

55 M. A. Fortuna, R. Ortega, and J. Bascompte, The web of life. Preprint arXiv:1403.2575 (2014).

56 J.-G. Young, plant-pollinator-inference (2021), URL https://doi.org/10.5281/zenodo.4759159.

